# Transcriptional and clonal characterization of cytotoxic CD8^+^ T cells in crescentic glomerulonephritis

**DOI:** 10.1101/2022.08.31.506004

**Authors:** Yu Zhao, Anne Mueller, Hakan Cicek, Hans-Joachim Paust, Amirrtavarshni Sivayoganathan, Alexandra Linke, Claudia Wegscheid, Thorsten Wiech, Tobias B. Huber, Catherine Meyer-Schwesinger, Stefan Bonn, Ulf Panzer, Gisa Tiegs, Christian F. Krebs, Katrin Neumann

## Abstract

Crescentic glomerulonephritis (cGN), most often caused by anti-neutrophil cytoplasmic autoantibody (ANCA)-associated vasculitis, is an aggressive form of immune-mediated kidney disease and represents an important cause of end-stage renal failure. Although it is known that T cells infiltrate the kidney in cGN, their precise role in autoimmune kidney disease remains to be fully elucidated. By performing single-cell analysis, we identified activated, clonally expanded CD8^+^ T cells with a cytotoxic gene expression profile in the kidneys of patients with ANCA-associated cGN. Using an experimental model of cGN, we demonstrated that clonally expanded murine CD8^+^ T cells highly expressed the cytotoxic molecule granzyme B. Moreover, lack of CD8^+^ T cells or granzyme B resulted in an ameliorated course of cGN. This was associated with reduced cleaved caspase-3 induction in renal tissue cells. Our data indicate that clonally expanded cytotoxic CD8^+^ T cells have a previously unrecognized pathogenic function in aggravating immune-mediated kidney disease.

## Introduction

Glomerulonephritis (GN) comprises a group of immune-mediated kidney diseases driven by inappropriately regulated cellular and humoral immune responses. Renal involvement of anti-neutrophil cytoplasmic autoantibody (ANCA)-associated vasculitis (AAV) results in crescentic GN (cGN) characterized by glomerular crescent formation and necrosis. This can lead to rapid decline in kidney function and end-stage renal disease, which is associated with high morbidity and mortality^1,2^.

Genome-wide association studies conducted in patients with AAV identified several genetic variants in HLA-DP and HLA-DQ regions, that associate with disease susceptibility^3,4^, highlighting that T cells contribute to disease pathology. Indeed, persistent inflammatory CD4^+^ T cell activation promotes autoimmunity in AAV^5,6,7^. In ANCA-associated cGN, CD4^+^ T cells recognizing the autoantigen myeloperoxidase (MPO) have been reported to mediate renal injury^8,9^. Elevated numbers of Th1 and Th17 cells have been described in blood^10,11,12^ and kidney^13^, indicating that both drive autoimmunity in ANCA-associated cGN. Although CD8^+^ T cells also infiltrate the kidney^14,15^, their functional role in autoimmune kidney disease is less clear. In experimental anti-MPO GN, MPO-specific CD8^+^ T cells were shown to mediate glomerular injury if MPO was planted in glomeruli^16^. Transcriptome profiling of blood CD8^+^ T cells in AAV revealed that CD8^+^ T cells with an activated, non-exhausted profile positively correlate with a poor prognosis^17^, suggesting their pathogenicity in AAV. However, phenotype and function of T cells depend on the organ in which they reside, and analyses done with blood cells might not reflect the situation in the inflamed kidney. To understand T cell-mediated immunity, it is imperative to perform a functional profiling of T cells from the kidneys of patients with cGN.

The mechanisms of T cell activation are still not well understood in autoimmune kidney disease. T cell receptors (TCRs) are responsible for antigen recognition and subsequent T cell activation and clonal expansion. T cell clones are defined as a group of T cells arising from one single naïve T cell, which all share the same TCR. To ensure a high TCR diversity, TCRs are generated via V(D)J recombination. The hypervariable complementarity-determining region 3 (CDR3) is the most common V region of the TCR and interacts with antigenic peptides thereby determining the antigen specificity of a TCR^18^. Sequencing of TCRs at single cell levels allows to unravel size and diversity of TCR repertoires^19,20^, which may help to understand T cell-mediated immune responses in autoimmune kidney disease.

Here, we performed a deep analysis of transcriptional profiles and TCR sequences of T cells from the inflamed kidney in ANCA-associated cGN. Cytotoxic T lymphocytes (CTLs) revealed the highest clonal expansion and displayed an activated profile. Expanded renal CD8^+^ T cells in experimental cGN showed high expression of the cytotoxic molecule granzyme B. Our functional analyses demonstrated that lack of CD8^+^ T cells or granzyme B ameliorated renal damage. We therefore described mechanisms of CD8^+^ T cell-mediated injury in cGN.

## Results

### Clonal expansion of cytotoxic CD8^+^ T cells in ANCA-associated cGN

To understand the functional heterogeneity of T cell subsets in autoimmune kidney disease, we performed single-cell RNA sequencing (scRNA-seq) in combination with cellular indexing of transcriptomes and epitopes by sequencing (CITE-seq) and single-cell TCR sequencing (scTCR-seq) of CD3^+^ T cells isolated from renal biopsies and blood of patients with ANCA-associated cGN (Fig. 1a). As control, we used healthy kidney tissue from patients undergoing tumour nephrectomy. After quality control and filtering (Supplemental Fig. 1), we included 8.810 renal T cells from nine patients with ANCA-associated cGN and 12.336 renal T cells from five control patients in our analysis. We applied principal component analysis on variably expressed genes and proteins across all CD3^+^ T cells to generate uniform manifold approximation and projection (UMAP). We identified 11 main clusters including effector CD4^+^ and CD8^+^ T cell subsets, granzyme B^+^ (*GZMB*) CTLs, *FOXP3*^+^ regulatory T cells (Tregs), natural cytotoxicity triggering receptor 1^+^ (*NCR1*)/T cell receptor delta variable 2^+^ (*TRDV2*) NKT cells/γδ T cells and T cell receptor alpha variable 1-2 (*TRAV1-2*) MAIT cells (Fig. 1b, c, Supplemental Fig. 2a). All T cell subsets were distributed across each sample (Supplemental Fig. 2b).

**Fig. 1:**
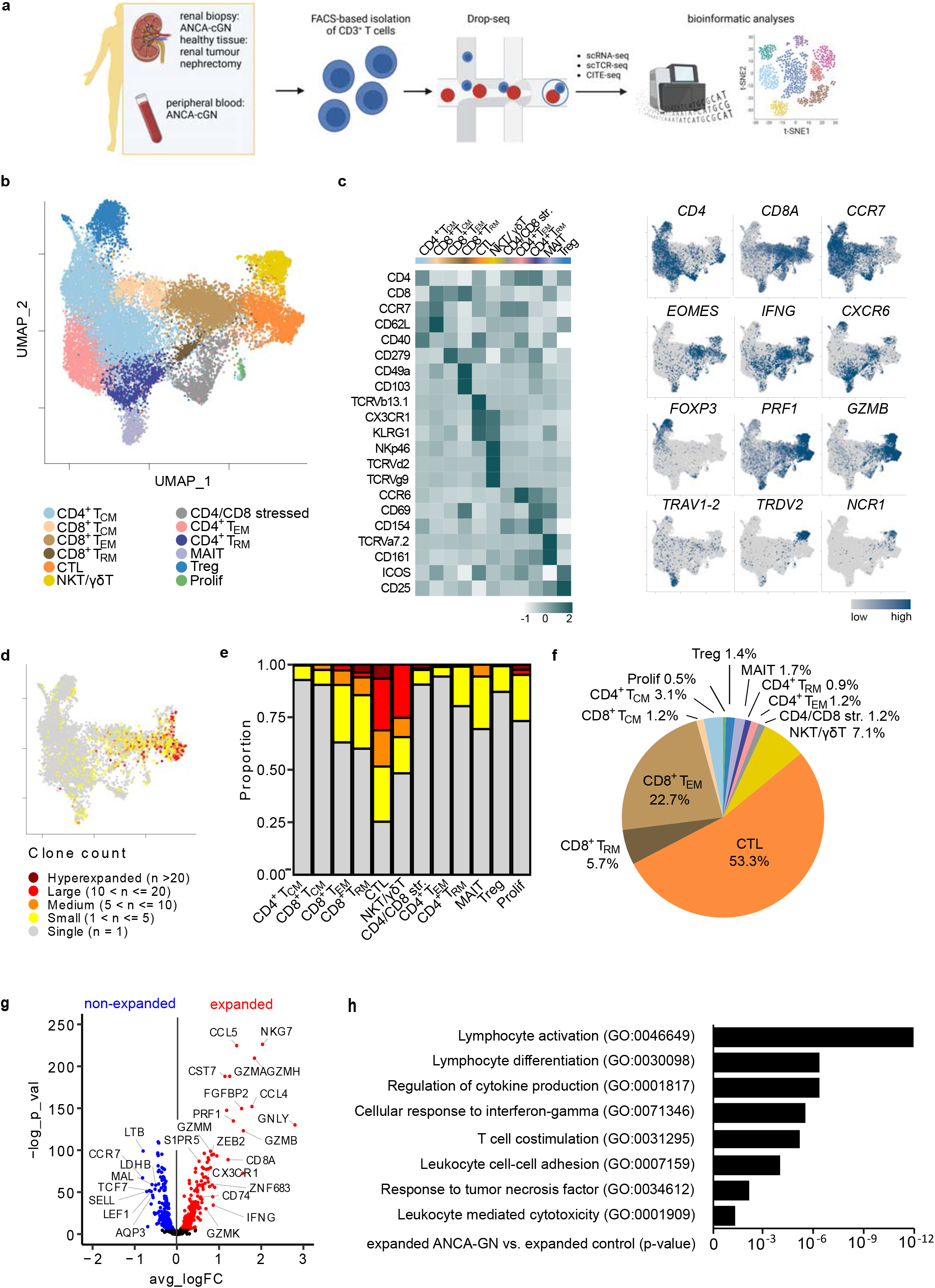
Clonally expanded renal T cell subsets in ANCA-associated cGN. **a** Schematic overview of the study design. FACS-isolated CD3^+^ T cells from kidney biopsies and peripheral blood of patients with ANCA-associated cGN and healthy renal tissue of patients undergoing tumour nephrectomy were subjected to scRNA-seq, CITE-seq and scTCR-seq analyses. **b** UMAP plot shows clustering of renal T cell subsets from ANCA-associated cGN and control patients. **c** Heat map and UMAP plots show subset-defining protein and gene expression. **d** UMAP plot shows overlay of renal T cell subsets and clonally expanded T cells from patients with ANCA-associated cGN. **e** Proportion of single to hyperexpanded renal T cells is depicted for each T cell subset. **f** Frequencies of expanded renal T cells within each T cell subset are shown. **g** Volcano plot depicts the most upregulated genes in expanded renal T cells compared to non-expanded T cells. **h** GO analysis of upregulated pathways in clonally expanded renal T cells from patients with ANCA-associated cGN in comparison to control patients.

To assess clonal expansion of T cells in ANCA-associated cGN, we quantified similar T cell clones based on the TCR and the nucleotide sequence of the CDR3 region and overlaid the scTCR-seq information on top of the defined T cell subsets. We determined clonally expanded renal T cells in patients with ANCA-associated cGN (Fig. 1d, Supplemental Fig. 2c). The vast majority of them were CD8^+^ T cells and to a much lesser extent NKT/γδ T cells. The CTL cluster showed the largest proportion of T cell clones and constituted the majority of highly expanded T cells (Fig. 1e, f). Comparative transcriptome analysis revealed strongly upregulated expression of *GZMB, GZMA*, granulysin (*GNLY*), perforin 1 (*PRF1*) and *CX3CR1* in expanded compared to non-expanded T cells, demonstrating the accumulation of cytotoxic CD8^+^ T cell clones in ANCA-associated cGN (Fig. 1g). Gene ontology (GO) pathway analysis revealed that T cell clones from patients with ANCA-associated cGN were highly activated in comparison to the control samples, in which clonally expanded T cells were also predominantly found in the CTL cluster (Supplemental Fig. 2d, e). However, pathways involved in T cell activation and differentiation were enriched in clonally expanded T cells in ANCA-associated cGN (Fig. 1h). These data indicate that activated and cytotoxic CD8^+^ T cell clones are present in the kidneys of patients with ANCA-associated cGN and might contribute to disease pathology.

### Systemic distribution of T cell clonotypes in ANCA-associated cGN

Since cGN is part of a systemic inflammation in AAV, we compared the TCR repertoires of renal and peripheral blood CD3^+^ T cells obtained from the same patients. Therefore, 10.504 blood T cells were included in our analysis. Similar to the kidney, the majority of clonally expanded T cells in the blood were detected within CD8^+^ T cells (Fig. 2a-d) and were characterized by upregulated expression of genes associated with cytotoxicity (Fig. 2e). By analyzing the most expanded T cell clones, we showed that they are present in both kidney and blood of the same patient with ANCA-associated cGN (Fig. 2f), indicating their systemic distribution, which mirrors the systemic vasculitis.

**Fig. 2:**
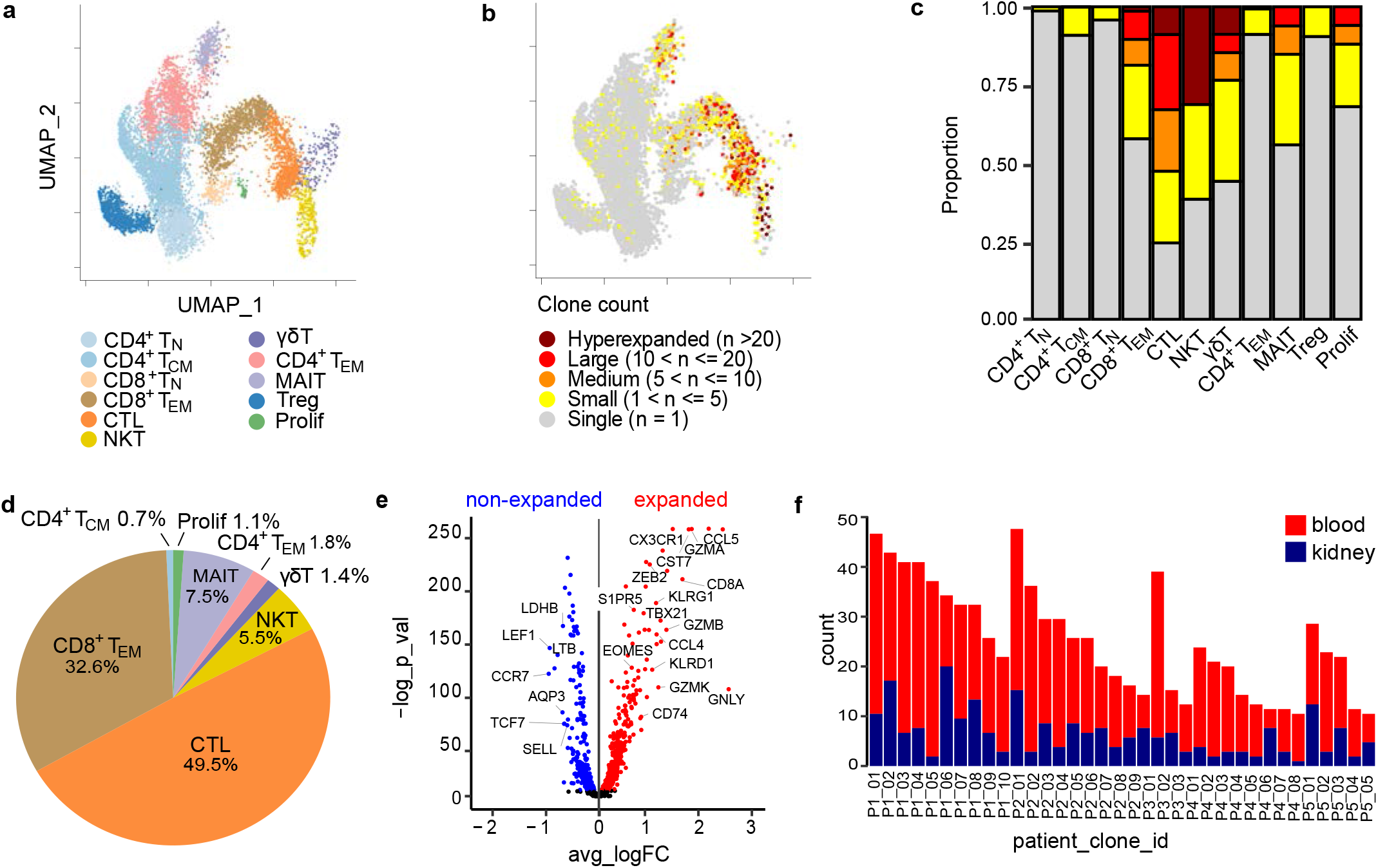
Clonally expanded peripheral blood T cell subsets in ANCA-associated cGN. **a** UMAP plot shows clustering of peripheral blood T cell subsets in patients with ANCA-associated cGN. **b** UMAP plot shows overlay of blood T cell subsets and clonally expanded T cells. **c** Proportion of single to hyperexpanded blood T cells is depicted for each T cell subset. **d** Frequencies of expanded blood T cells within each T cell subset are shown. **e** Numbers of the most expanded T cell clones in kidney and blood of the same patient are shown.

### Different effector CD8^+^ T cell subsets in ANCA-associated cGN

After detection of cytotoxic CD8^+^ T cell clones, we addressed whether pathogenic CD8^+^ T cells promote disease pathology of ANCA-associated cGN. To get an overview about functional CD8^+^ T cell subsets, we performed a phenotype analysis of CD8^+^ T cells in ANCA-associated cGN. Based on described subset signatures^21,22,23,24^, we identified three subsets of renal CD8^+^ T cells: cytotoxic effector CD8^+^ T cells (T_Eff_; *GZMB*^+^ CD69^-^ CD27^-^ CD62L^low^ CCR7^-^), in which most of the clonally expanded CD8^+^ T cells were present, effector memory/central memory CD8^+^ T cells (T_EM/CM_; *GZMB*^-^ CD69^low^ CD27^+^ CD62L^int^ CCR7^+^) and tissue-resident memory CD8^+^ T cells (T_RM_; *GZMB*^-^ CD69^+^ CD27^-^ CD62L^-^ CCR7^low^). In blood, we detected naïve CD8^+^ T cells (T_N_), while the T_RM_ subset was absent (Fig. 3a, Supplemental Fig. 3a, b). Canonical marker expression analysis confirmed our subset definition: CD8^+^ T_Eff_ expressed genes associated with activation and cytotoxicity^25,26,27,28^ (*KLRG1, GZMB, GZMH, GNLY, PRF1, CX3CR1*) and tissue egress^29,30^ (*S1PR1*/*5, KLF2*/*3*). CD8^+^ T_EM/CM_ specifically expressed *GZMK* as well as *CXCR4* and *CRTAM*, markers linked with homeostasis, migration and retention of memory T cells^31,32,33^. CD8^+^ T_RM_ expressed *ZNF683*, which directs the transcriptional program of tissue residency, and related genes^21,34,35^ like *CXCR6, ITGA1* and *XCL1*. In contrast, genes associated with tissue egress were less expressed (Fig. 3b). Differentially expressed gene (DEG) analysis revealed subset-specific gene expression profiles in kidney (Figure 3c) and blood (Supplemental Fig. 3c). Venn diagrams showed less overlap of upregulated genes in the different CD8^+^ T cell subsets from kidney and blood. GO pathway analysis highlighted compartment- and subset-specific differences. For example, cytotoxicity-associated pathways were overrepresented in renal CD8^+^ T_Eff_, whereas migration and cell activation processes were upregulated in blood CD8^+^ T_Eff_ (Fig. 3d). Transcription factor analysis also showed compartment- and subset-specific variations in CD8^+^ T cells in ANCA-associated cGN (Supplemental Fig. 3d, e). We did the same set of analyses with a previously published data set from patients with ANCA-associated cGN^36^, thereby confirming our CD8^+^ T cell subset definition (Supplemental Fig. 4).

**Fig. 3:**
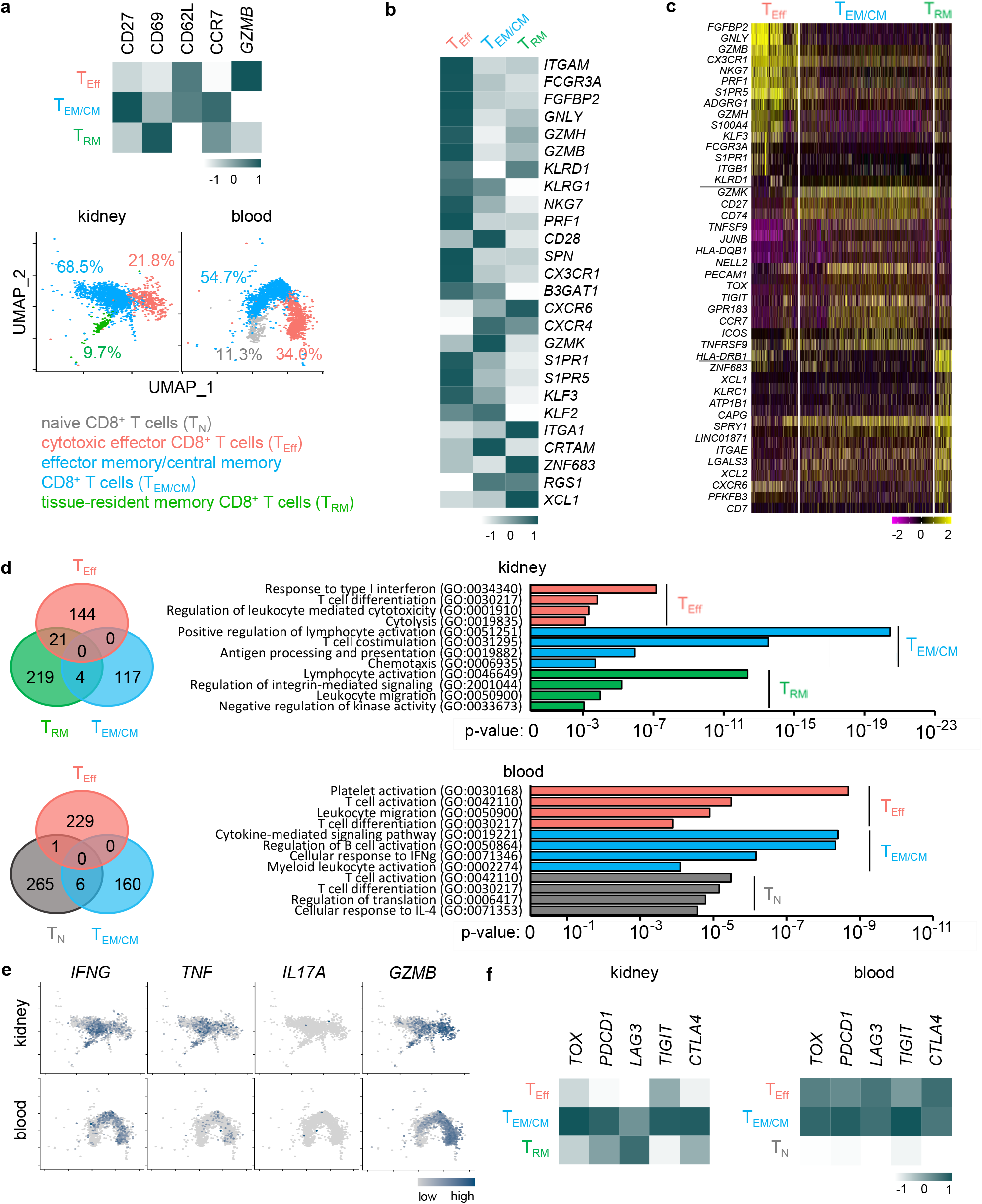
Gene expression profile of CD8^+^ T cell subsets in ANCA-associated cGN. **a** Heat map of CD8^+^ T cell subset-defining protein and gene expression in the kidney. UMAP plots show clustering and percentages of CD8^+^ T cell subsets from kidney and blood. **b** Heat map of subset-defining gene expression profiles of each renal CD8^+^ T cell subset. Heat maps show the average normalized expression of signature genes in the individual subsets. **c** Heat map depicts the 15 most upregulated genes for each CD8^+^ T cell subset in the kidney. **d** Overlap of upregulated genes in indicated CD8^+^ T cell subsets is represented by Venn diagram. GO analysis of upregulated pathways in CD8^+^ T cell subsets from kidney and blood. **e** UMAP plots of cytokine and *GZMB* gene expression in renal and blood CD8^+^ T cells subsets. **f** Heat maps of marker genes for T cell exhaustion in renal and blood CD8^+^ T cells subsets.

Besides cytotoxicity, CD8^+^ T cells also drive immunity by production of inflammatory cytokines. Therefore, we analyzed cytokine expression and identified high expression of *IFNG* and *TNF* in CD8^+^ T cells from patients with ANCA-associated cGN. Although IL-17A-producing CD8^+^ T cells have been described in other autoimmune diseases^37^, we could not detect expression of this cytokine by CD8^+^ T cells in patients with ANCA-associated cGN. By comparing the different CD8^+^ T cell subsets, we found that *IFNG* and *TNF* were particularly expressed by renal CD8^+^ T_EM/CM_ and CD8^+^ T_RM_, while the *GZMB*-expressing CD8^+^ T_Eff_ subset showed less inflammatory cytokine expression (Fig. 3e).

Prolonged T cell activation in the context of chronic inflammation is a key driver of T cell dysfunction, termed exhaustion^38^. To assess CD8^+^ T cell exhaustion, we analyzed expression of genes associated with immune regulation and exhaustion and found that in patients with ANCA-associated cGN, particularly the renal CD8^+^ T_EM/CM_ subset expressed *TOX, PDCD1, LAG3, TIGIT* and *CTLA4*, while these genes were less expressed in CD8^+^ T_Eff_. In blood, these genes were expressed by T_Eff_ and T_EM/CM_ but not T_N_ (Fig. 3f, Supplemental Fig. 3f). These data indicate a particular role of CD8^+^ T_Eff_ in disease pathology by expression of cytotoxic mediators in the kidney.

### Clonal expansion of cytotoxic CD8^+^ T cells in murine cGN

To address the functional role of CD8^+^ T cells in cGN, we used a well-established murine model of experimental cGN. We induced cGN in wild-type (WT) mice, isolated CD3^+^ T cells from the inflamed kidneys and subjected these cells to scRNA-seq in combination with scTCR-seq. After quality control and filtering, we included 6119 renal T cells in the analyses (Fig. 4a). In mice, NKT cells and γδ T cells clustered separately, while a distinct cluster of MAIT cells was not present. There was a cluster of interferon-responsive T cells, which expressed interferon-stimulated genes (ISG). Comparable to ANCA-associated cGN, we detected a CTL cluster characterized by expression of *Gzmb* (Fig. 4b, Supplemental Fig. 5a). Interestingly, we also determined clonally expanded T cells in murine cGN. The distribution of clones across T cell subsets in experimental cGN was similar to that observed in the kidneys of patients with ANCA-associated cGN with the highest proportion of clonally expanded T cells in the CTL cluster (Fig. 4c-e). We determined upregulated expression of genes associated with cytotoxicity (*Gzmb, Gzma* and *Prf1*) and *Cd8a* in expanded versus non-expanded T cells (Fig. 4f). Thus, clonally expanded cytotoxic CD8^+^ T cells were also present in the kidneys of mice with cGN.

**Fig. 4:**
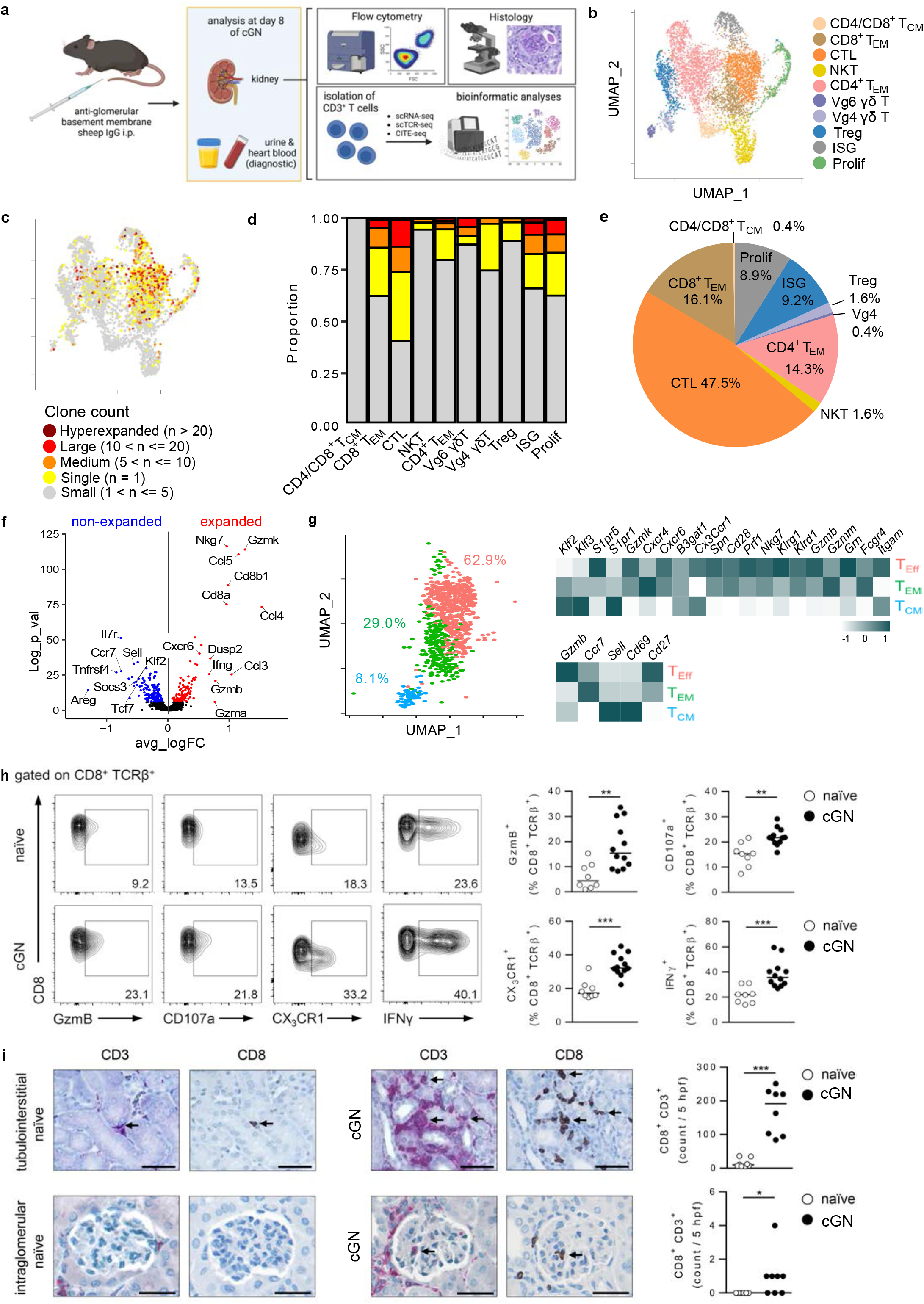
Clonal expansion and phenotype of CD8^+^ T cell subsets in murine cGN. Experimental cGN was induced in WT mice and kidneys were analyzed 8 days later. **a** Schematic overview of the study design. **b** UMAP plot shows clustering of renal T cell subsets in nephritic WT mice. **c** Overlay of renal T cell subsets and clonally expanded T cells is shown. **d** Proportion of single to hyperexpanded renal T cells is depicted for each T cell subset. **e** Frequencies of expanded renal T cells within each T cell subset are shown. **f** Volcano plot depicts the most upregulated genes in expanded renal T cells compared to non-expanded T cells. **g** Heat maps and UMAP plot show CD8^+^ T cell subset-defining gene expression. **h** Protein expression of renal CD8^+^ T cells was analyzed in naïve and nephritic WT mice. **i** Serial kidney sections of naïve and nephritic WT mice were stained with anti-CD3 or anti-CD8 antibodies. Bars represent 50 μm (tubulointerstitial) or 25 μm (intraglomerular). Numbers of tubulointerstitial and intraglomerular CD8^+^ CD3^+^ T cells were counted. Arrows mark CD8^+^ T cells. Representative dot plots and medians of two experiments are shown. *p< 0.05; **p< 0.01; ***p< 0.001.

A detailed analysis of all renal CD8^+^ T cells in experimental cGN revealed that the majority of them were CD8^+^ T_Eff_. We also detected CD8^+^ T_EM_ and T_CM_, while CD8^+^ T_RM_ could not be identified. Particularly CD8^+^ T_Eff_ and to a lesser extent CD8^+^ T_EM_ were characterized by expression of genes associated with cytotoxicity. In murine cGN, CD8^+^ T_Eff_ also expressed *Gzmk* and *Cxcr6* (Fig. 4g, Supplemental Fig. 5b-d). We further determined expression of *Ifng* and *Tnf* by CD8^+^ T_Eff_ and T_EM_, while *Il17a* was not expressed. In murine cGN, CD8^+^ T_Eff_ showed the strongest expression of immune regulatory genes (Supplemental Fig. 5e).

By analyzing protein expression, we demonstrated elevated frequencies of renal CD8^+^ T cells expressing GzmB, the degranulation marker CD107a^39^ and IFN-γ in nephritic compared to naïve WT mice. The frequency of CX_3_CR1^+^ CD8^+^ T cells was also increased in experimental cGN, a chemokine receptor, which has been associated with cytotoxic CD8^+^ T cells^28^ (Fig. 4h). We used viSNE analysis to identify distinct CD8^+^ T cell clusters based on protein expression. The analysis revealed distinct clusters of degranulated (GzmB^low/-^CD107a^+^) and not degranulated (GzmB^+^CD107a^-^) CD8^+^ T cells. IFN-γ^+^ CD8^+^ T cells clustered separately with small subsets also expressing GzmB or CD107a (Supplemental Fig. 5f). To assess localization of renal CD8^+^ T cells, we stained for CD8 and CD3 in serial kidney sections of naïve and nephritic WT mice. As expected, in naïve mice, the number of CD8^+^ CD3^+^ T cells was low, while we determined increased CD8^+^ T cell numbers in the inflamed kidney. The majority of these CD8^+^ T cells accumulated in the tubulointerstitium (Fig. 4i), reflecting the situation in the kidneys of patients with ANCA-associated cGN (Supplemental Fig. 5g). Thus, inflammatory CD8^+^ T cells infiltrate the kidney in murine cGN and accumulate in the tubulointerstitium where they are found in close contact to tubular epithelial cells.

### Lack of CD8^+^ T cells ameliorates experimental cGN

To study the functional role of CD8^+^ T cells in cGN, we induced experimental cGN in *Cd8a*^-/-^ mice, which lack CD8^+^ T cells (Fig. 5a), while CD4^+^ T cells were not affected (Supplemental Fig. 6a). Pathologically, CD8^+^ T cell-deficient mice displayed reduced crescent formation, proteinuria and blood urea nitrogen (BUN) levels compared to WT mice with cGN (Fig. 5b, c). To understand the mechanisms behind CD8^+^ T cell pathogenicity in cGN, we investigated the immune cell composition in *Cd8a*^-/-^ and WT mice. This analysis showed no differences in the frequencies of renal CD4^+^ T cells, CD11c^+^ dendritic cells (DCs) and Foxp3^+^ Tregs in naïve WT and *Cd8a*^-/-^ mice (Fig. 5d-f). In the spleen, the frequency of CD4^+^ T cells was elevated in naïve *Cd8a*^-/-^ mice, whereas no alterations were found in DCs and Tregs (Supplemental Fig. 6b-d). In nephritic mice, renal CD4^+^ T cells showed an elevated expression of the co-inhibitory receptor PD-1 and the cytokines IFN-γ and IL-17A in the absence of CD8^+^ T cells (Fig. 5d). We further determined a reduced frequency of Tregs and DCs in the kidneys of nephritic *Cd8a*^-/-^ compared to WT mice (Fig. 5e, f). These differences were not observed in splenic immune cell populations (Supplemental Fig. 6b-d). We also stained for F4/80 in renal tissue and demonstrated a reduced number of F4/80^+^ macrophages in nephritic *Cd8a*^-/-^ mice (Fig. 5g). Thus, *Cd8a*^-/-^ mice develop less severe cGN, which is associated with reduced renal infiltration of macrophages.

**Fig. 5:**
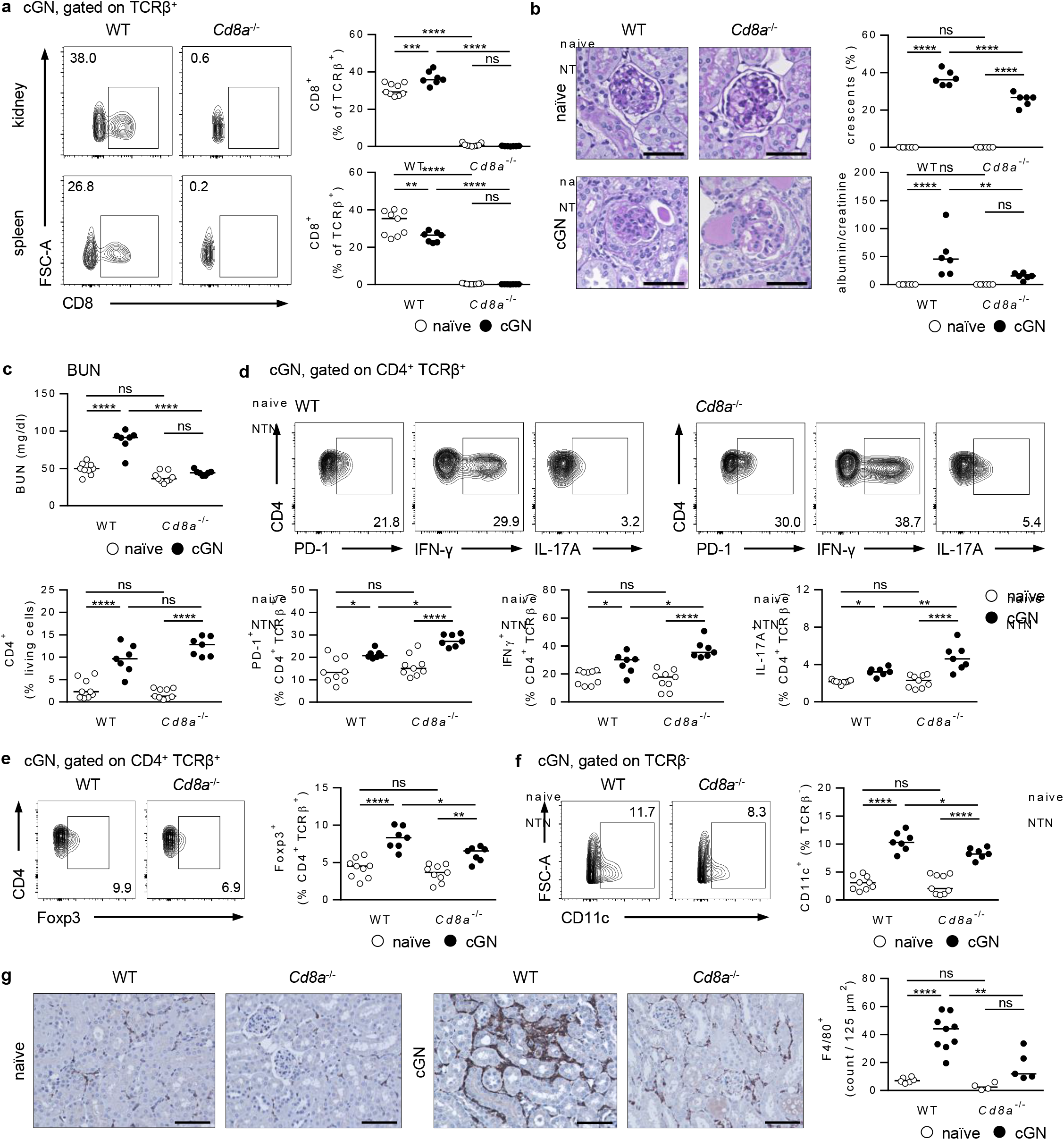
Disease pathology of experimental cGN in *Cd8a*^-/-^ mice. Crescentic GN was induced in *Cd8a*^-/-^ and WT mice, that were analyzed 8 days later. **a** Frequencies of renal and splenic CD8^+^ T cells were analyzed in naïve and nephritic *Cd8a*^-/-^ and WT mice. **b** Glomerular crescent formation was quantified in PAS-stained kidney sections. Bars represent 50 μm. Proteinuria was assessed by determining the albumin-to-creatinine ratio in urine. **c** BUN levels were determined in serum. Frequencies and phenotypes of renal (**d**) CD4^+^ T cells, (**e**) Foxp3^+^ Tregs and (**f**) CD11c^+^ DCs were analyzed in naïve and nephritic *Cd8a*^-/-^ and WT mice. **g** Kidney sections of naïve and nephritic *Cd8a*^-/-^ and WT mice were stained with an anti-F4/80 antibody. Numbers of F4/80^+^ cells were counted. Bars represent 50 μm. Representative dot plots and medians of one out of two experiments are shown. *p< 0.05; **p< 0.01; ***p< 0.001; ****p< 0.0001, ns: not significant.

### Less severe renal pathology in the absence of GzmB

After demonstrating the presence of GzmB-expressing CD8^+^ T cells in human and murine cGN, including a clonally expanded fraction, we focused on the role of this cytotoxic molecule in disease pathology. One mechanism by which GzmB induces apoptotic cell death in target cells is via activation of procaspase-3 through proteolytic cleavage^40^. Therefore, we stained for cleaved caspase-3, the active form of this enzyme, in renal tissue. In murine cGN, we detected an elevated number of cleaved caspase-3^+^ cells in nephritic compared to naïve WT mice. Expression of cleaved caspase-3 was predominantly observed in tubular epithelial cells. Induction of cleaved caspase-3 in tubular epithelial cells and intraglomerular cells was reduced in *GzmB*^-/-^ mice (Fig. 6a). In these mice, CD8^+^ T cells did not express GzmB (Fig. 6b). As a consequence, *GzmB*^-/-^ mice showed reduced crescent formation and BUN levels compared to nephritic WT mice, while proteinuria was not altered (Fig. 6c, d). There were no differences in frequencies and phenotypes of renal and splenic CD8^+^ T cells, CD4^+^ T cells and Tregs in nephritic *GzmB*^-/-^ compared to WT mice. The frequency of DCs was decreased in kidney but not spleen (Fig. 6e, Supplemental Fig. 7a, b). Moreover, infiltration of F4/80^+^ macrophages was not reduced in the absence of GzmB (Fig. 6f). Thus, *GzmB*^-/-^ mice develop less severe cGN, which is associated with reduced induction of cleaved caspase-3 particularly in tubular epithelial cells. Interestingly, induction of cleaved caspase-3 in tubular epithelial cells and intraglomerular cells was also reduced in nephritic *Cd8a*^-/-^ mice (Fig. 6g), demonstrating the pro-apoptotic function of cytotoxic CD8^+^ T cells in murine cGN.

**Fig. 6:**
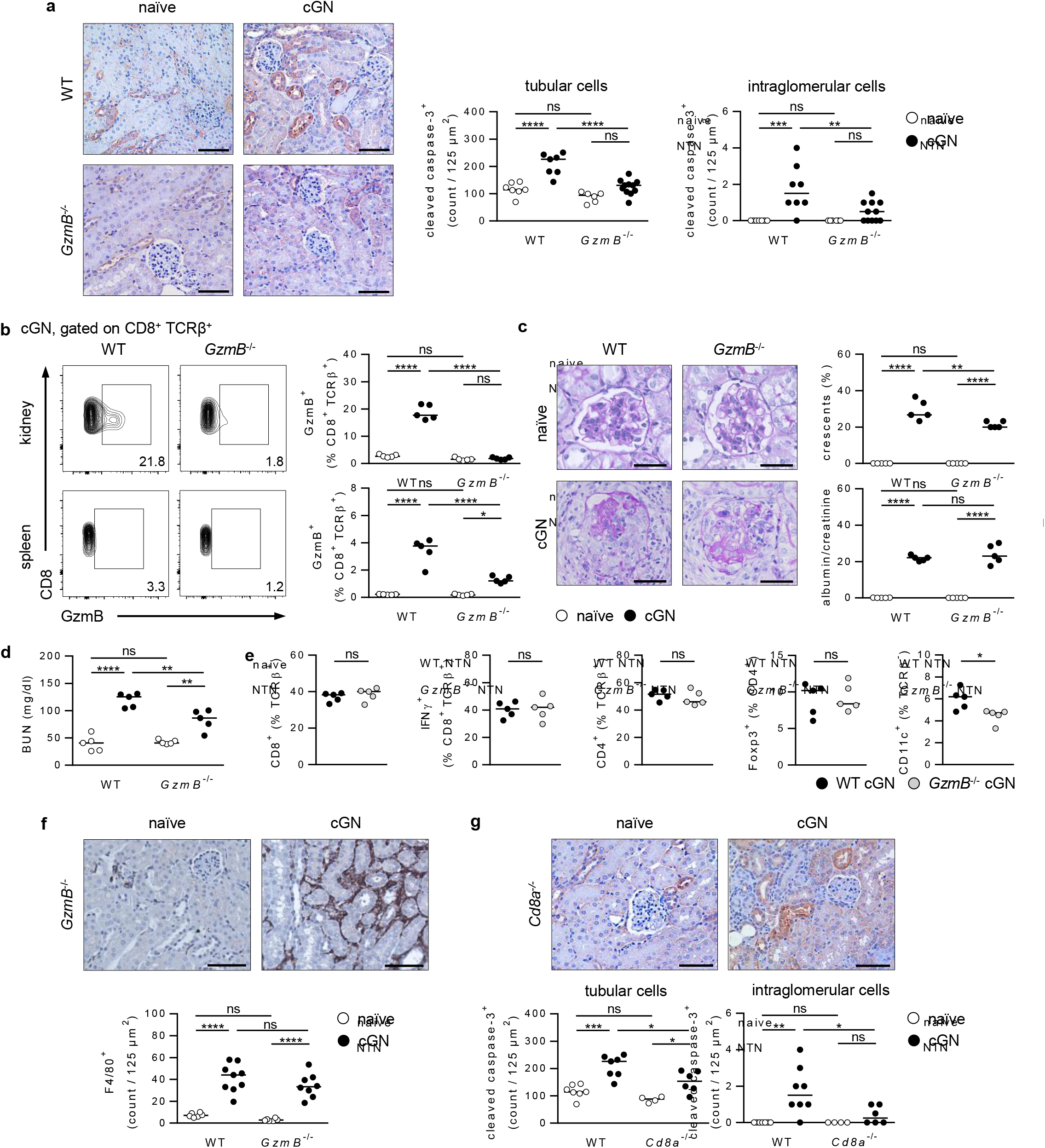
Disease pathology of experimental cGN in *GzmB*^-/-^ mice. **a** Crescentic GN was induced in *GzmB*^-/-^ and WT mice, that were analyzed 8 days later. Kidney sections were stained with an anti-cleaved caspase-3 antibody. Numbers of cleaved caspase-3^+^ cells were counted. Bars represent 50 μm. **b** GzmB expression in CD8^+^ T cells was analyzed. **c** Glomerular crescent formation was quantified in PAS-stained kidney sections. Bars represent 100 μm. Proteinuria was assessed by determining the albumin-to-creatinine ratio in urine. **d** BUN levels were determined in serum. **e** Frequencies of renal CD8^+^ T cells, Foxp3^+^ Tregs, CD4^+^ T cells and CD11c^+^ DCs were analyzed in nephritic *GzmB*^-/-^ and WT mice. **f** Kidney sections of naïve and nephritic *GzmB*^-/-^ mice were stained with an anti-F4/80 antibody. Numbers of F4/80^+^ cells were counted. Bars represent 50 μm. **g** Kidney sections of naïve and nephritic *Cd8a*^-/-^ were stained with an anti-cleaved caspase-3 antibody. Numbers of cleaved caspase-3^+^ cells were counted. Bars represent 50 μm. Representative dot plots and medians of one out of two experiments are shown. *p< 0.05; **p< 0.01; ***p< 0.001; ****p< 0.0001, ns: not significant.

## Discussion

Crescentic GN is a live-threatening disease triggered by so far poorly defined mechanisms. This results in lack of specific therapeutic strategies not relying on overall immunosuppression. Thus, identifying novel immunopathogenic mechanisms in cGN is of high clinical relevance. In this study, we showed for the first time the presence of clonally expanded renal cytotoxic CD8^+^ T cells in ANCA-associated cGN and experimental cGN. We further described GzmB-induced tubular apoptosis as a novel mechanism of CD8^+^ T cell-mediated immunity in cGN.

There is increasing evidence that CD8^+^ T cells contribute to the induction and progression of autoimmune diseases^41^. We identified activated, cytotoxic CD8^+^ T cell clones in kidney and blood of patients with ANCA-associated cGN, indicating that these cells circulate throughout the body and may impact renal and also systemic inflammation in AAV. It needs to be further addressed whether clonally expanded CD8^+^ T cells recognize autoantigens in ANCA-associated cGN, as it has been shown for MPO-specific CD4^+^ T cells^8,9^. There is considerable evidence that infections in genetically susceptible individuals initiate immune responses leading to autoimmunity, as shown in AAV^42,43^. It might be conceivable that CD8^+^ T cells are activated during such an infection, clonally expand and persist over time. The longevity of CD8^+^ T cell clones was shown in lupus nephritis, where they persisted for years, probably being responsible for disease progression^44^.

By performing immune cell profiling in ANCA-associated cGN, we identified functionally distinct CD8^+^ T cell subsets in the inflamed kidney. While the CD8^+^ T_Eff_ subset was characterized by high expression of *GZMB* and *PRF1*, the memory CD8^+^ T_EM/CM_ subset specifically expressed *GZMK*. CD8^+^ T cell subsets expressing either GzmB or GzmK have also been described in lupus nephritis^45^. Analyses of cytotoxic granule profiles in human CD8^+^ T cells revealed distinct granzyme expression pattern associated with the stage of differentiation and cytotoxic activity^46,47,48^. Cytotoxic CD8^+^ T cell function was strongly correlated with expression of perforin and GzmB^49,50^, whereas GzmK expression has been described for memory CD8^+^ T cells with less cytotoxic capacity^51,52^. This is in line with our data demonstrating low *GZMB* and *PRF1* expression in the memory CD8^+^ T cell subset. A recent study determined GzmK-expressing CD8^+^ T cells in human inflammatory diseases including lupus nephritis. The authors also showed a low cytotoxic potential of GzmK^+^ CD8^+^ T cells and highlighted them as major cytokine producers^53^. Interestingly, we found a similar correlation in patients with ANCA-associated cGN, since the inflammatory cytokines *IFNG* and *TNF* were predominantly expressed by *GZMK*^+^ CD8^+^ T_EM/CM_ and not by *GZMB*^+^ CD8^+^ T_Eff_.

Recently, we have defined T_RM_ with a Th1 and Th17 cell signature in the kidneys of patients with ANCA-associated cGN and the number of total T_RM_ positively correlated with impaired renal function, demonstrating their potential pathogenicity in autoimmune kidney disease^36^. In the present study, we determined CD8^+^ T_RM_ in ANCA-associated cGN, that showed less expression of cytotoxicity-associated genes but expressed *IFNG* and *TNF* and therefore, might also contribute to disease pathology. That CD8^+^ T_RM_ can aggravate autoimmune kidney disease has been shown in lupus prone MRL-*lpr* mice, in which the number of CD8^+^ T_RM_ increased during the course of the disease and correlated with severity of lupus nephritis^54^. So far, little is known about mechanisms leading to the activation of CD8^+^ T_RM_ in autoimmune disease. Pathogens infecting the kidney were found to induce IL-17A-producing T_RM_17, which persisted in the kidney after pathogen clearance and aggravated murine cGN due to reactivation by local inflammatory cytokines^36^. CD8^+^ T_RM_ have also been shown to develop during infections and persist in peripheral tissue to provide rapid protection against reinfection^55^. Therefore, it might be possible that also CD8^+^ T_RM_ become reactivated in target tissues of autoimmunity. In line with this, cytokine-mediated activation of CXCR6^+^ CD8^+^ T_RM_ has been described in a mouse model of chronic liver disease^56^.

To assess mechanisms of CD8^+^ T cell-mediated injury in autoimmune kidney disease, we used a murine model of cGN. Immune cell profiling of murine renal T cells revealed that the highest number of clonally expanded T cells were again found in the cytotoxic CD8^+^ T_Eff_ subset. In contrast to patients with ANCA-associated cGN, we did not determine CD8^+^ T_RM_ in experimental cGN. This can be explained by the fact that the mouse model is a rather acute model of cGN and that young mice housed under specific pathogen-free conditions were used for the experiments, which prevented previous infections possibly leading to induction of CD8^+^ T_RM_. We did not observe such a strong difference in the gene expression profiles of CD8^+^ T_Eff_ and CD8^+^ T_EM_ as shown for ANCA-associated cGN. It rather seems that due to the more acute inflammation, we depicted CD8^+^ T cell transition states as we detected expression of *Gzmk* and *Cxcr6* in CD8^+^ T_Eff_, two genes associated with memory T cells, whereas CD8^+^ T_EM_ still expressed *Gzmb* and *Prf1*. Cytokine gene expression analysis revealed no significant difference in CD8^+^ T_Eff_ and CD8^+^ T_EM_ but by analyzing protein expression, we found that also in murine cGN, GzmB- and IFN-γ-expressing CD8^+^ T cells mainly cluster separately.

We showed that *Cd8a*^-/-^ mice developed less severe renal pathology in experimental cGN, demonstrating their pathogenicity in this model. We made sure that the improved pathology was not a result of impaired CD4^+^ T cell activation in the absence of CD8^+^ T cells. Previous studies have found that IFN-γ^+^ Th1 cells promote renal tissue damage via recruitment of inflammatory macrophages^9,57,58^. Since CD8^+^ T cells produced IFN-γ in cGN, we analyzed renal macrophage infiltration in nephritic *Cd8a*^-/-^ mice. Indeed, the number of macrophages was reduced in these mice, indicating inflammation-induced recruitment of macrophages as one mechanism by which CD8^+^ T cells promote kidney injury in cGN.

In line with our findings, CD8^+^ T cells have been shown predominantly in the tubulointerstitial compartment of kidneys of patients with ANCA-associated cGN^14,15^. This localization was also observed in murine cGN, where CD8^+^ T cells were identified in close proximity to tubular epithelial cells. These cells displayed an elevated induction of cleaved caspase-3, the active form of procaspase-3, critically involved in apoptotic cell death. From all granzymes, only GzmB induces proteolytic activation of procaspase-3 in target cells and thus, uses the caspase cascade for induction of apoptosis^40^. Since nephritic *GzmB*^-/-^ mice showed reduced induction of cleaved caspase-3 in tubular epithelial cells, this demonstrates that GzmB promotes tubular apoptosis in murine cGN. Lack of GzmB improved pathogenesis of cGN although not as pronounced as in the absence of CD8^+^ T cells. That is probably because CD8^+^ T cells expressing IFN-γ were still present in *GzmB*^-/-^ mice, thus, renal macrophage infiltration was not altered in the absence of GzmB. Importantly, we also detected decreased induction of cleaved caspase-3 in tubular epithelial cells from nephritic *Cd8a*^-/-^ mice, demonstrating that in murine cGN, cytotoxic CD8^+^ T cells promote apoptosis in the tubular system by release of GzmB. This finding is supported by a recent study, in which we showed that also in lupus nephritis, lack of CD8^+^ T cells resulted in reduced induction of tubular apoptosis^59^.

In summary, this study revealed functionally distinct renal CD8^+^ T cell subsets in ANCA-associated cGN, that can contribute to disease pathology by GzmB-induced tubular apoptosis and cytokine-induced inflammatory macrophage recruitment. Moreover, highly activated and clonally expanded cytotoxic CD8^+^ T cell clones, which are present in kidney and blood, might trigger aggravation and progression of both renal and systemic inflammation in AAV. This study further highlights GzmB as a novel therapeutic target in autoimmune kidney disease, which needs to be further investigated in future trails.

## Methods

### Human samples

Kidney biopsies were analyzed from patients with ANCA-associated cGN included in the Hamburg GN Registry^36^. Matched blood samples from the same patients were used for the analyses. As controls, healthy renal tissue from patients undergoing tumour nephrectomy or zero biopsies of renal transplants were used. These studies were approved by the Ethik-Kommission der Ärztekammer Hamburg and were conducted in accordance with the ethical principles stated in the Declaration of Helsinki (Registration numbers PV 5026 and PV 5822). Informed consent was obtained from all participating patients.

### Animals

C57BL/6J (WT), B6.129S2-*Cd8a*^tm1Mak/^J (*Cd8a*^-/-^), and B6.129S2-*Gzmb*^tm1Ley/^J (*GzmB*^-/-^) mice were bred in the animal facility of the University Medical Center Hamburg-Eppendorf (UKE; Hamburg, Germany). Mice were bred according to the Federation of European Laboratory Animal Science Association guidelines and maintained under specific-pathogen-free conditions. All mouse experiments were approved by the Behörde für Justiz und Verbraucherschutz (Hamburg, Germany) and carried out according to the current existing guidelines on mouse experimentation.

### Animal treatment

To induce experimental cGN (nephrotoxic nephritis), 8-10 weeks old WT, *Cd8a*^-/-^, and *GzmB*^-/-^ mice received an intraperitoneal injection of sheep serum (2.5 mg/g body weight) containing antibodies directed against the murine glomerular basement membrane and were analyzed 8 days later^13^.

### Urine analysis

One day before final analysis, mice were housed in metabolic cages for urine collection. Albumin levels were determined by ELISA (Mice-Albumin Kit; Bethyl, Montgomery, TX). Creatinine levels were measured with the COBAS INTEGRA Creatinine Jaffé Gen.2 Kit (Roche Diagnostics, Indianapolis, IN). To assess severity of proteinuria, the albumin-to-creatinine ratio was calculated.

### Serum analysis

Serum BUN levels were measured with the COBAS INTEGRA BUN Kit (Roche Diagnostics, Indianapolis, IN) 8 days after induction of cGN.

### Histological analyses

Histology was performed as previously described^60^. In brief, periodic acid-Schiff (PAS) staining was done with 1.5 μm paraffin-embedded kidney sections to determine crescent formation, which was assessed in 30 glomeruli/mouse in a blinded fashion. CD8 and CD3 staining was performed in serial sections to identify CD8^+^ CD3^+^ T cells in renal tissue from human and mice. 2 μm paraffin-embedded, serial human kidney sections were stained with anti-CD3 (LN10) or anti-CD8 (4B11; both Leica Biosystems, Chicago, IL) antibodies. The ZytoChem-Plus AP Polymer-Kit (Zytomed Systems) was used for antigen detection. For murine analysis, 2 μm paraffin-embedded kidney sections were stained with anti-CD3 (polyclonal, Agilent Technologies, Santa Clara, CA) or anti-CD8 (D4W2Z; Cell Signaling Technology, Danvers, MA) antibodies. A horseradish peroxidase (HRP)-conjugated anti-rabbit antibody (ThermoFisher Scientific, Waltham, MA) was used as secondary antibody. The DAB+ Substrate Chromogen System (Agilent Technologies) was used for antigen detection. For antigen detection, New Fuchsin (Merck, Darmstadt, Germany) and the ZytoChem-Plus AP Polymer-Kit (Zytomed Systems, Berlin, Germany) were used. To stain cleaved caspase-3, 2 μm paraffin-embedded kidney sections were incubated with an anti-cleaved caspase-3 antibody (ThermoFisher Scientific). After incubation with goat anti-rabbit-HRP secondary antibody (ThermoFisher Scientific), kidney sections were stained with DAB+ substrate (Agilent Technologies). To stain F4/80, 2 μm paraffin-embedded kidney sections were incubated with an anti-F4/80 antibody (CI:A3-1; Bio-Rad, Feldkirchen, Germany). After incubation with biotinylated goat anti-rat IgG (Abcam, Cambridge, UK), kidney sections were stained with VECTASTAIN ABC-HRP Kit, Peroxidase (Biozol) and subsequent DAB+ substrate. For analysis, kidney sections were scanned using the Zeiss Axioscan 7 (Carl Zeiss, Jena, Germany) and analyzed by ZEN lite software (Carl Zeiss). To assess CD8^+^ T cells, five corresponding high power fields (hpf) were randomly defined in the cortices of each serial kidney section. Overlapping CD3 and CD8 staining marked CD8^+^ T cells. To assess cleaved caspase-3^+^ and F4/80^+^ cells, positive cells were counted in three random areas (each 125 μm^2^) located in renal cortex, from which the mean value was calculated.

### Isolation of renal leukocytes and splenocytes

Kidneys and spleens were harvested from healthy and nephritic mice 8 days after induction of cGN. Splenic tissue was passed through 70 μm nylon meshes prior to erythrocyte lysis using NH_4_Cl. Kidneys were finely minced and digested for 40 min at 37°C with 0.4 mg/ml collagenase D (Roche, Mannheim, Germany) and 0.01 mg/ml DNase I (Roche). Single cell suspensions were achieved by using the gentleMACS Dissociator (Miltenyi Biotec). To enrich renal leukocytes, cell suspensions were subjected to a density gradient centrifugation using a 37% Percoll solution (GE Healthcare Life Sciences). Erythrocytes were lysed with NH_4_Cl.

### Flow cytometry

Renal leukocytes and splenocytes were stimulated with phorbol myristate acetate (10 ng/ml) and ionomycin (250 ng/ml) for 4 hours with the addition of brefeldin A (1 μg/ml; all Sigma Aldrich) and monensin (BioLegend) after 30 minutes. The anti-CD107a antibody (1D4B; FITC; BioLegend) was added to the medium. Cells were incubated with anti-CD16/32 antibody solution (93; BioLegend, San Diego, CA) to prevent unspecific antibody binding. LIVE/DEAD Fixable Staining Kits (ThermoFisher Scientific) were used to exclude dead cells. Cells were stained with fluorochrome-labeled antibodies specific to TCRβ (H57-597; FITC), CD8a (53-6.7; BV 785), CX_3_CR1 (SA011F11; BV 711), CD11c (N418; PE-Dazzle 594), CD4 (RM4-5; BV 711) and PD-1 (J43; FITC; all BioLegend). After fixation and permeabilization (Foxp3 Transcription Factor Staining Buffer Set; ThermoFisher Scientific), cells were stained with antibodies specific to GzmB (GB11; Pacific Blue), IFN-γ (XMG1.2; PE-Texas Red), IL-17A (TC11-18H10.1; PerCP) and Foxp3 (MF-14; AF 647; all BioLegend). Analyses were done using a BD LSRFortessa Cell Analyzer (BD Biosciences).

### Single cell RNA sequencing

To perform scRNA-seq, single cell suspensions were obtained from samples of human and mouse kidneys. FACS-sorted CD3^+^ T cells were subjected to droplet-based single cell analysis (Chromium, 10x Genomics, Pleasanton, CA) using the Chromium Single Cell 5′ Reagent Kits v2 chemistry according to the manufacturer’s instructions. TCR and gene expression libraries were sequenced on a NovaSeq 6000 System (150 cycles paired-end; Illumina, San Diego, CA^61^).

### Alignment, quality control and pre-processing of scRNA-seq data

The Cell Ranger software (version 5.0.1; 10X Genomics) was used to demultiplex cellular barcodes and map reads to the reference genomes (refdata-cellranger-hg19-1.2.0 (human) or refdata-gex-mm10-2020-A (mouse)). The CITE-seq antibody and barcode information was included in a feature reference CSV file and passed to the cellranger count command. As output, we obtained feature-barcode matrices containing gene expression counts alongside CITE-seq counts for each cell barcode. The feature-barcode matrices were further processed by the R package Seurat (version 4.0.2). For quality control, we filtered out all cells in which less than 500 genes or more than 6000 genes were detected. Low-quality cells with more than 10% mitochondrial genes of all detected genes were also removed from analysis (Supplemental Fig. 1). We used the LogNormalize method in Seurat to normalize the scRNA-seq and CITE-seq counts for all cells that have passed the quality control. We used Seurat demultiplexing function HTODemux to demultiplex the hash-tag samples.

### Dimensionality reduction and clustering

To reduce batch effects, we applied the integration method implemented in Seurat (function FindIntegrationAnchors and IntegrateData, dims = 1:30). The integrated matrix was then scaled by ScaleData function (default parameters). In order to reduce the dimensionality of the integrated data, principal component analysis was performed on the scaled data (function RunPCA, npcs = 30). 30 principal components were determined by using the ElbowPlot function and used to compute the KNN graph based on the euclidean distance (function FindNeighbors), which then generated cell clusters using function FindClusters. The resolution was chosen based on the cell cluster marker genes in the downstream analysis. Uniform Manifold Approximation and Projection (UMAP) was used to visualize clustering results. The top differential expressed genes in each cluster were found using the FindAllMarkers function (min.pct = 0.1) and ran Wilcoxon rank sum tests. The differential expression between clusters were calculated by FindMarkers function (min.pct = 0.1), which also ran Wilcoxon rank sum tests.

### Processing of scTCR-seq data and integration

scTCR-seq data for each sample were assembled by the Cell Ranger software (version 5.0.1) with the command cellranger vdj using the reference genome (refdata-cellranger-vdj-GRCh38-alts-ensembl-3.1.0). For each sample, Cell Ranger generated an output file, filtered_contig_annotations.csv, containing TCR-α chain and TCR-β chain CDR3 nucleotide sequences for single cells, that were identified by barcodes. The R package scRepertoire (version 1.5.2^62^) was used to further combine the contig_annotation data of different samples to a single list object (function combineTCR). The combined TCR contig list file was then integrated with the corresponding Seurat object of the scRNA-seq data using the function combineExpression (cloneCall = “gene+nt”). Only the cells with both TCR and scRNA-seq data were kept for downstream clonotype analysis. The clonotype was defined according to the genes comprising the TCR and the nucleotide sequence of the CDR3 region. The frequency of each clonotype in each patient was then calculated as clone count. The clone counts were added to metadata of the single-cell matrices.

### Pathway and transcription factor analysis

Gene ontology enrichment analyses have been performed with EnrichrR (version 1.0.^63^). We used the databases “GO_Biological_Process_2018”, “GO_Cellular_Component_2018” and “GO_Molecular_ Function_2018”. For transcription factor pathway activities, we used the tool DoRothEA^64^.

### Statistical analysis

Data were analyzed using the GraphPad Prism software (GraphPad software, San Diego, CA). Statistical comparison between two individual groups was carried out using the two-tailed Mann-Whitney U test. In case of three or more groups, the one-way ANOVA with Tukey’s multiple comparisons test was used. Data were presented as medians. A p value of less than 0.05 was considered statistically significant with the following ranges *p< 0.05, **p< 0.01, ***p< 0.001, and ****p< 0.0001.

## Acknowledgment

The authors acknowledge the excellent technical assistance of Alina Borchers, Elena Tasika, Carsten Rothkegel and Sonja Wulf.

## Author contributions

Y.Z. analyzed data, wrote and revised the manuscript; A.M. performed experiments, analyzed data, wrote and revised the manuscript; H.C. performed experiments, analyzed data; H.-J.P. performed experiments; A.S. performed experiments; A.L. performed experiments, analyzed data; C.W. performed experiments; T.W. renal histology, revised the manuscript; T.B.H. conception and design of research; C.M.-S. renal histology, revised the manuscript; S.B. supervised the data analysis and edited the manuscript; U.P. conception and design of research, revised the manuscript; G.T. conception and design of research, revised the manuscript; C.F.K. conception and design of research, revised and edited the manuscript; K.N. conception and design of research, performed experiments, analyzed data, wrote the manuscript.

## Data availability

Raw and processed data generated and analyzed during the current study will be provided via the NCBI GEO database (gene expression data are part of Paust et al. bioRxiv 2022).

## Abbreviations

AAV: anti-neutrophil cytoplasmic autoantibody-associated vasculitis
ANCA: anti-neutrophil cytoplasmic autoantibody
BUN: blood urea nitrogen
CDR3: complementarity-determining region 3
cGN: crescentic glomerulonephritis
CITE-seq: cellular indexing of transcriptomes and epitopes by sequencing
CTL: cytotoxic T lymphocyte
DC: dendritic cell
DEG: differentially expressed gene
GN: glomerulonephritis
GO: gene ontology
GzmB: granzyme B
MPO: myeloperoxidase
PAS: Periodic acid-Schiff
scRNA-seq: single-cell RNA sequencing
scTCR-seq: single-cell TCR sequencing
SLE: systemic lupus erythematosus
TCR: T cell receptor
T_Eff_: cytotoxic effector CD8^+^ T cell
T_EM/CM_: effector memory/central memory CD8^+^ T cell
T_N_: naive CD8^+^ T cells
Treg: regulatory T cell
T_RM_: tissue-resident memory CD8^+^ T cells
UMAP: uniform manifold approximation and projection
WT: wild-type

## Figure legends

**Supplemental Fig. 1:**
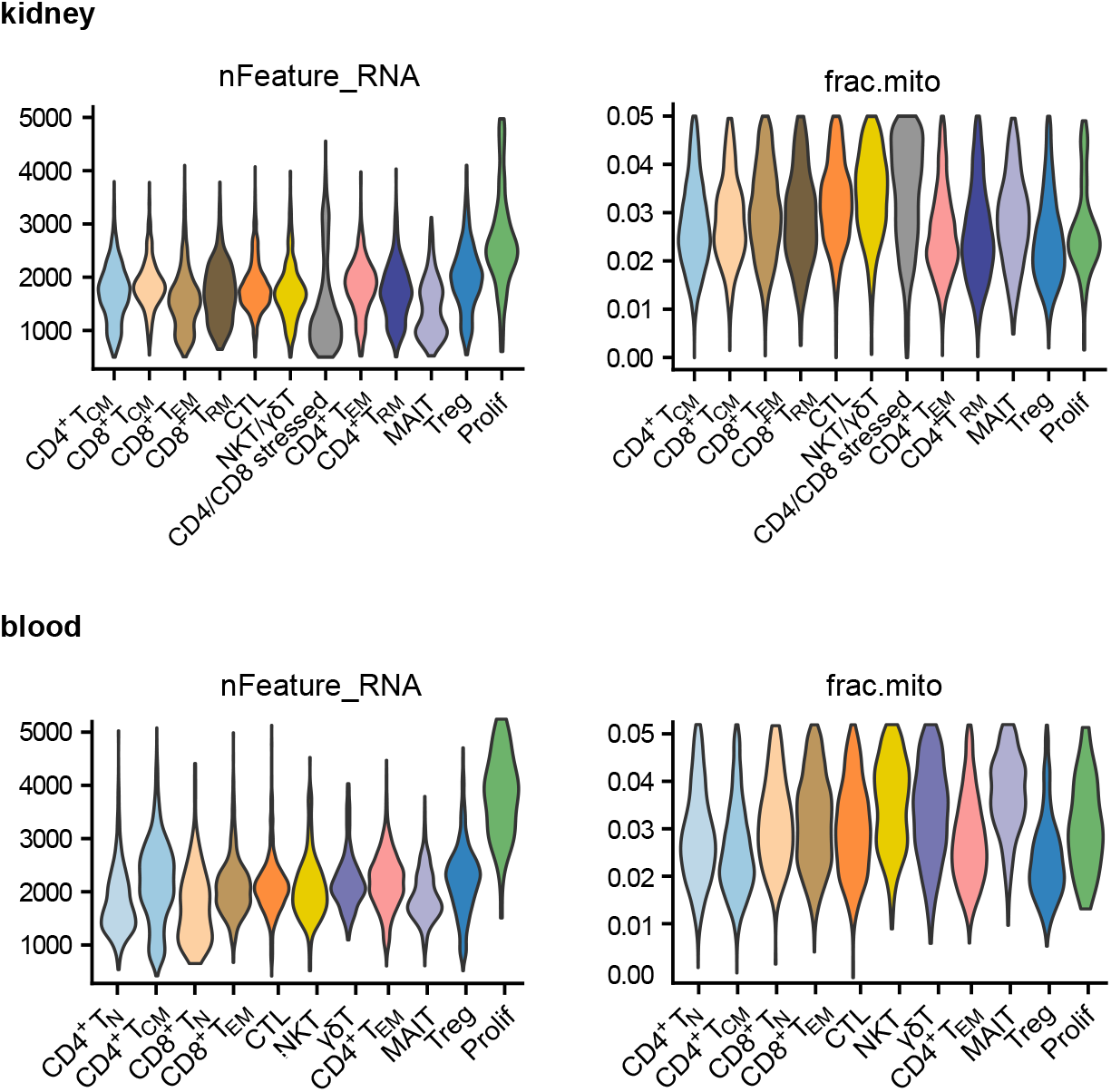
Quality control of the scRNA-seq data. CD3^+^ T cells isolated from kidney or blood of ANCA-cGN and control patients were pooled for the analyses. For quality control, all cells with less than 500 genes and more than 6000 genes were excluded. Low-quality cells with more than 10% mitochondrial genes of all detected genes were also excluded. nFeature_RNA represents the number of detected genes in each T-cell subset. Frac.mito represents the fraction of unique molecular identifiers (UMIs) in each T-cell subset aligned to mitochondrial genes.

**Supplemental Fig. 2:**
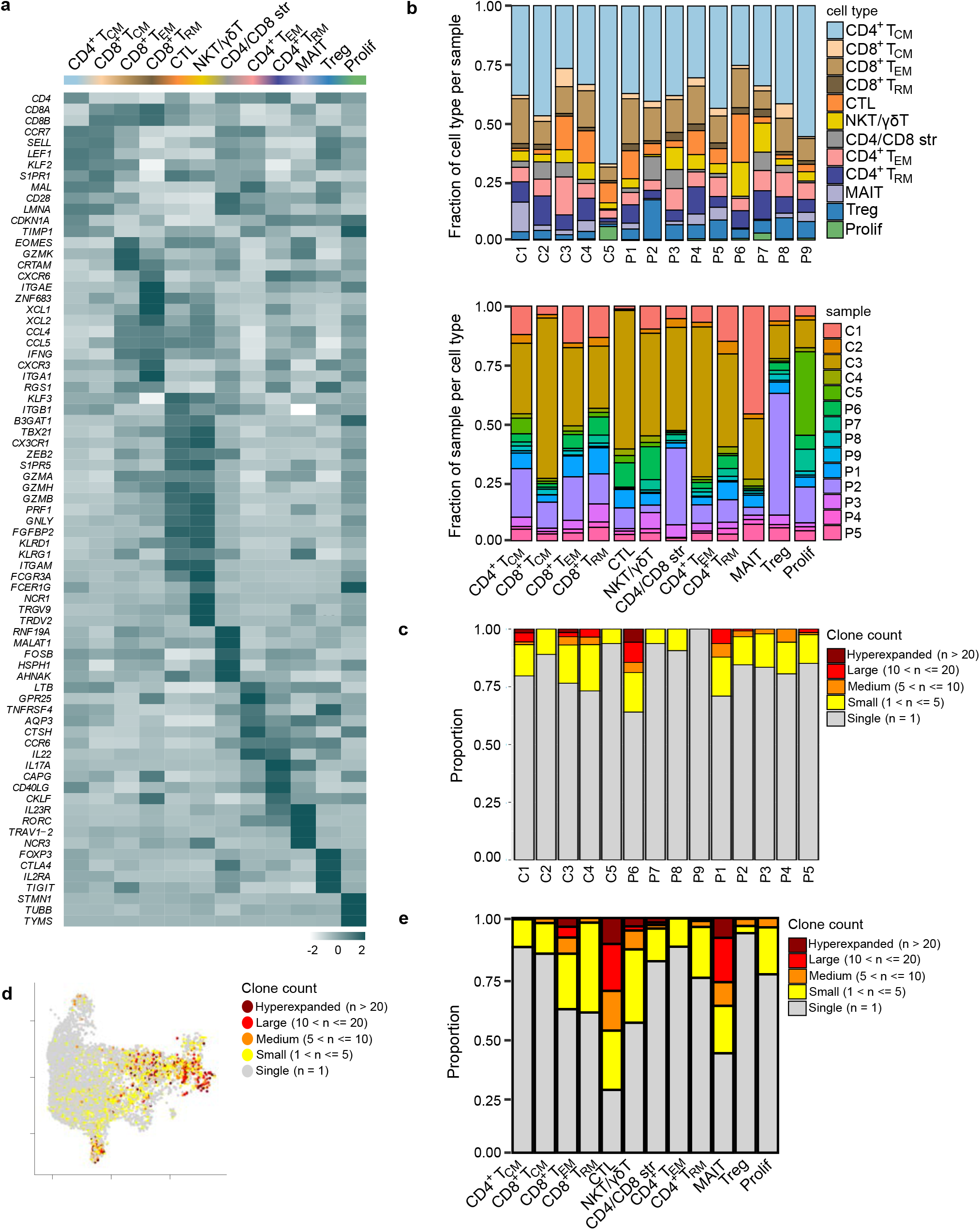
Immune cell profiling in ANCA-associated cGN and control patients. **a** Heat map shows gene expression profiles of the different T cell subsets in ANCA-associated cGN and control. **b** Distribution of the T cell subsets is depicted within each ANCA-associated cGN and control sample. **c** Distribution of clonally expanded T cells is shown in every ANCA-associated cGN sample. **d** UMAP plot shows overlay of renal T cell subsets and clonally expanded T cells from controls. **e** Proportion of single to hyperexpanded renal T cells is depicted for each T cell subset from controls.

**Supplemental Fig. 3:**
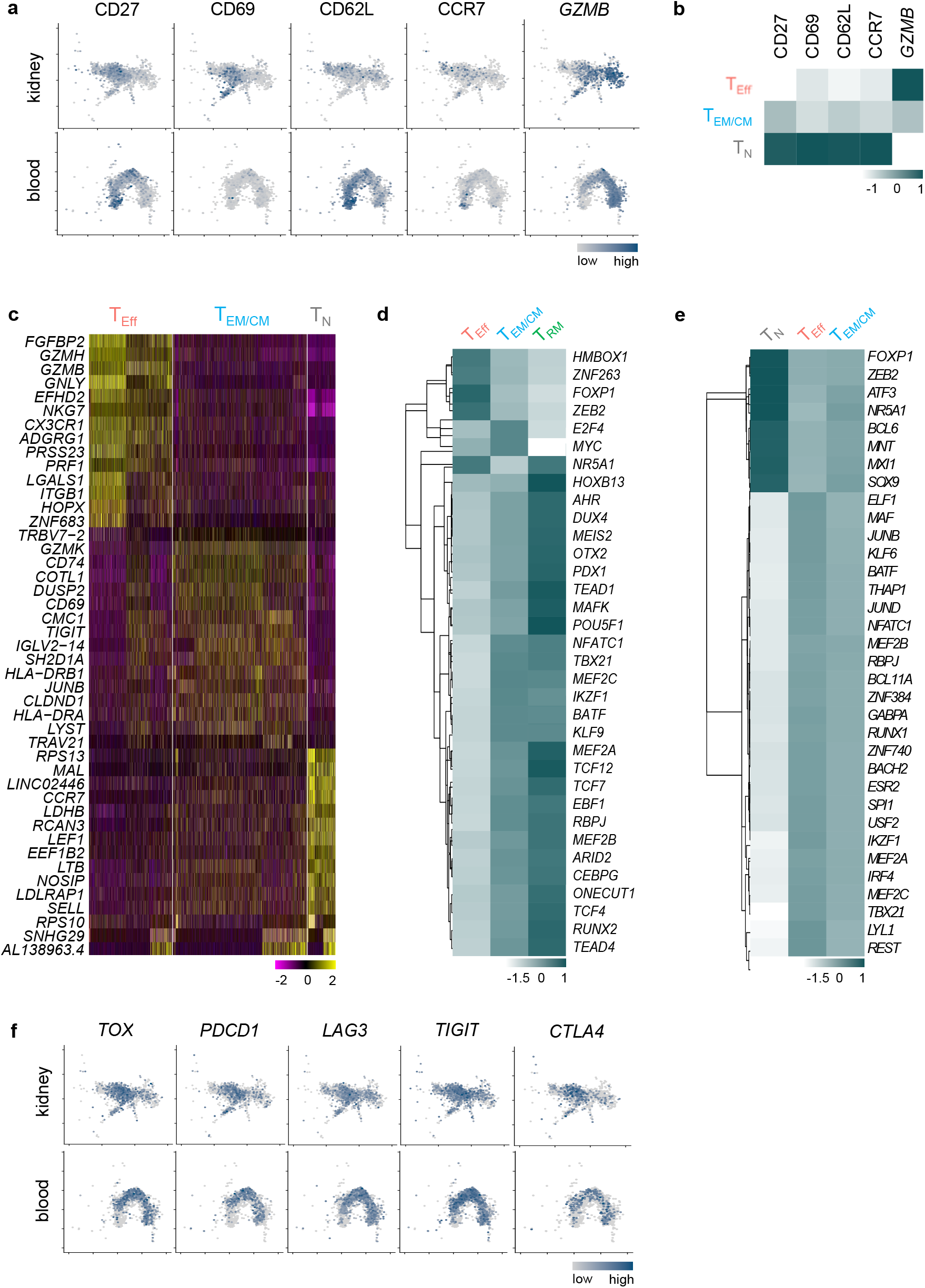
Gene and transcription factor profiles of CD8^+^ T cell subsets from patients with ANCA-associated cGN. **a** UMAP plots show marker gene expression defining CD8^+^ T cell subsets in kidney and blood. **b** Heat maps show CD8^+^ T cell subset-defining protein and gene expression in blood, c Heat map depicts the 15 most upregulated genes for each CD8^+^ T cell subset in blood. **d, e** Heat maps of transcription factor analysis in CD8^+^ T cells from **(d)** kidney and **(e)** blood. **f** UMAP plots show marker genes for immune regulation and T cell exhaustion in renal and blood CD8^+^ T cell subsets.

**Supplemental Fig. 4:**
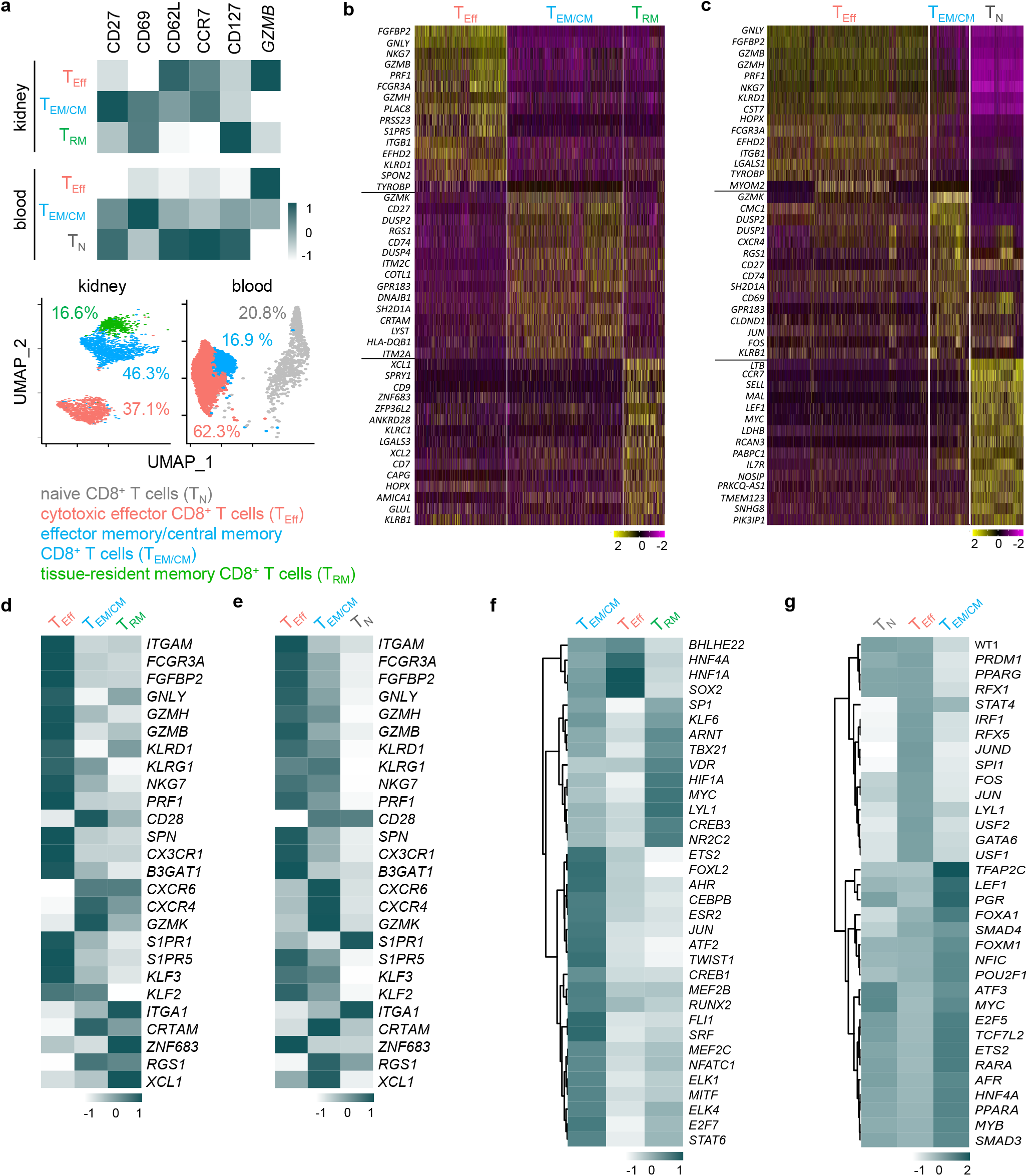
Gene expression profile of CD8^+^ T cell subsets from patients with ANCA-associated cGN. A previously published data set was used for the analyses (Krebs et al. Sci Immunol 2020) **a** Heat maps depict CD8^+^ T cell subset-defining protein and gene expression in kidney and blood. UMAP plots show clustering and percentages of CD8^+^ T cells subsets in kidney and blood. **b, c** Heat maps depict the 15 most upregulated genes for each CD8^+^ T cell subset in (**b**) kidney and (**c**) blood. **d, e** Heat map show subset-defining gene expression profiles of each CD8^+^ T cell subset in **(d)** kidney and **(e)** blood. **f, g** Heat maps show transcription factor analysis in CD8^+^ T cell subsets from (**f**) kidney and (**g**) blood.

**Supplemental Fig. 5:**
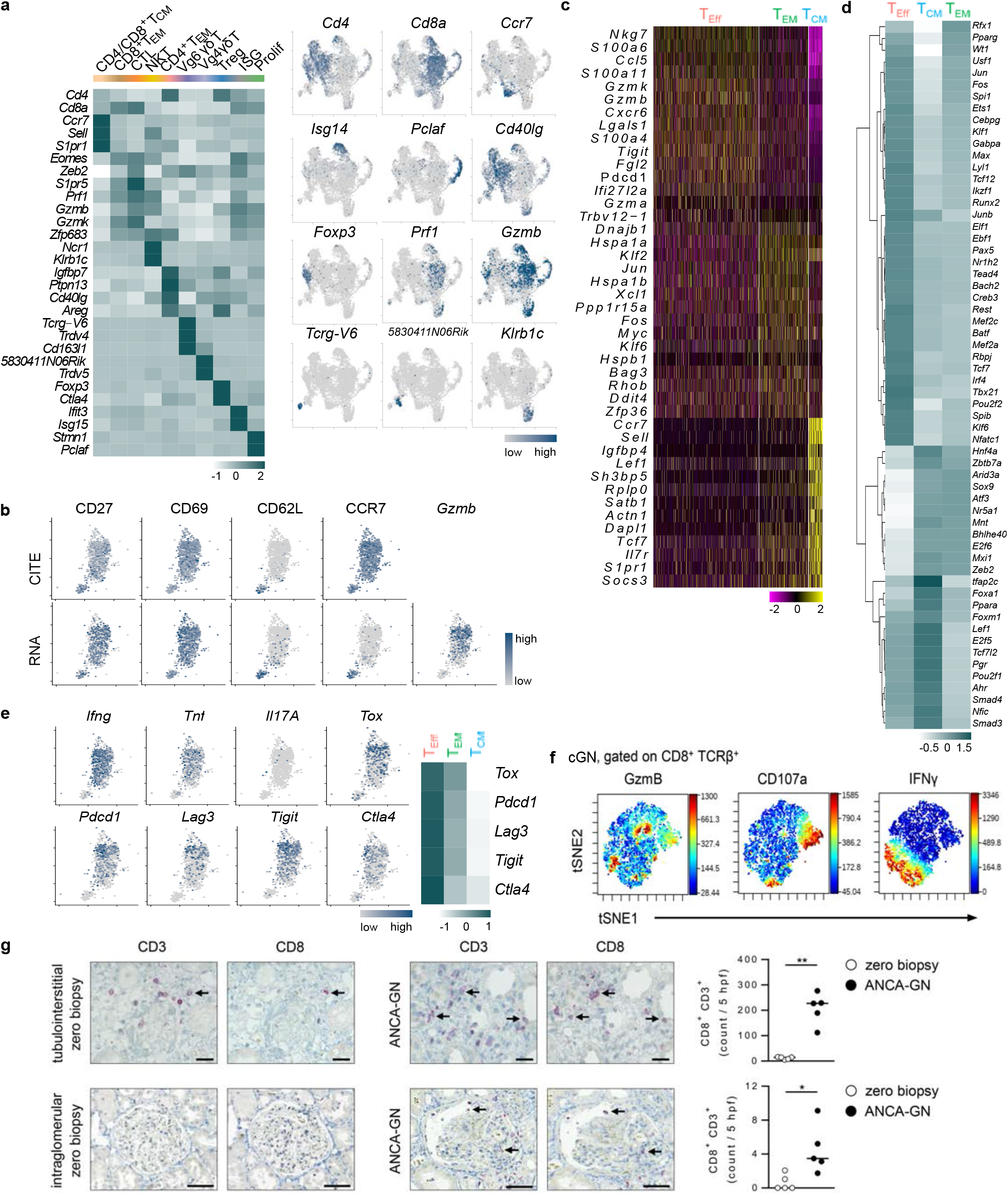
Phenotype of renal CD8^+^ T cells in experimental cGN. Crescentic GN was induced in WT mice, that were analyzed 8 days later, **a, b** Marker gene expression is shown for each T cell subset, **c, d** Heat maps depict (**c**) the most upregulated genes and (**d**) transcription factors for each CD8^+^ T cell subset, **e** Cytokine and immune regulatory gene expression, **f** ViSNE analysis of renal CD8^+^ T cells from nephritic WT mice comprising 3469 cells, **g** Serial kidney sections of patients with ANCA-associated cGN and zero biopsies were stained with anti-CD3 or anti-CD8 antibodies. Bars represent 25 pm (tubulointerstitial) or 50 pm (intra-glomerular). Arrows mark CD8^+^ T cells. *p< 0.05; **p< 0.01.

**Supplemental Fig. 6:**
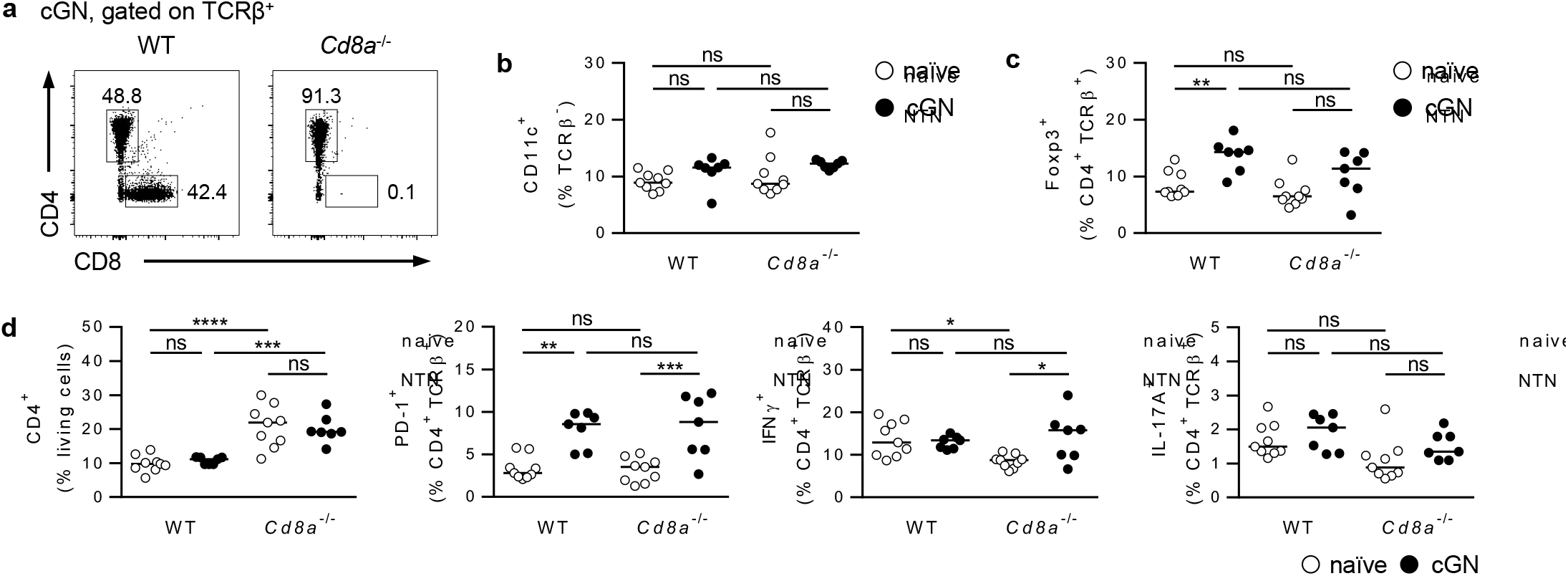
Splenic immune cell populations in *Cd8a*^-/-^ mice. Crescentic GN was induced in *Cd8a*^-/-^ and WT mice, which were analyzed 8 days later. **a** Frequencies of renal CD4^+^ and CD8^+^ T cells were shown in nephritic WT and *Cd8a*^-/-^ mice. Frequencies and phenotypes of splenic (**b**) CD11c^+^ DCs, (**c**) Foxp3^+^ Tregs and (**d**) CD4^+^ T cells were analyzed in naïve and nephritic *Cd8a*^-/-^ and WT mice. Medians of one out of two experiments are shown. *p< 0.05; **p< 0.01; ***p< 0.001; ns: not significant.

**Supplemental Fig. 7:**
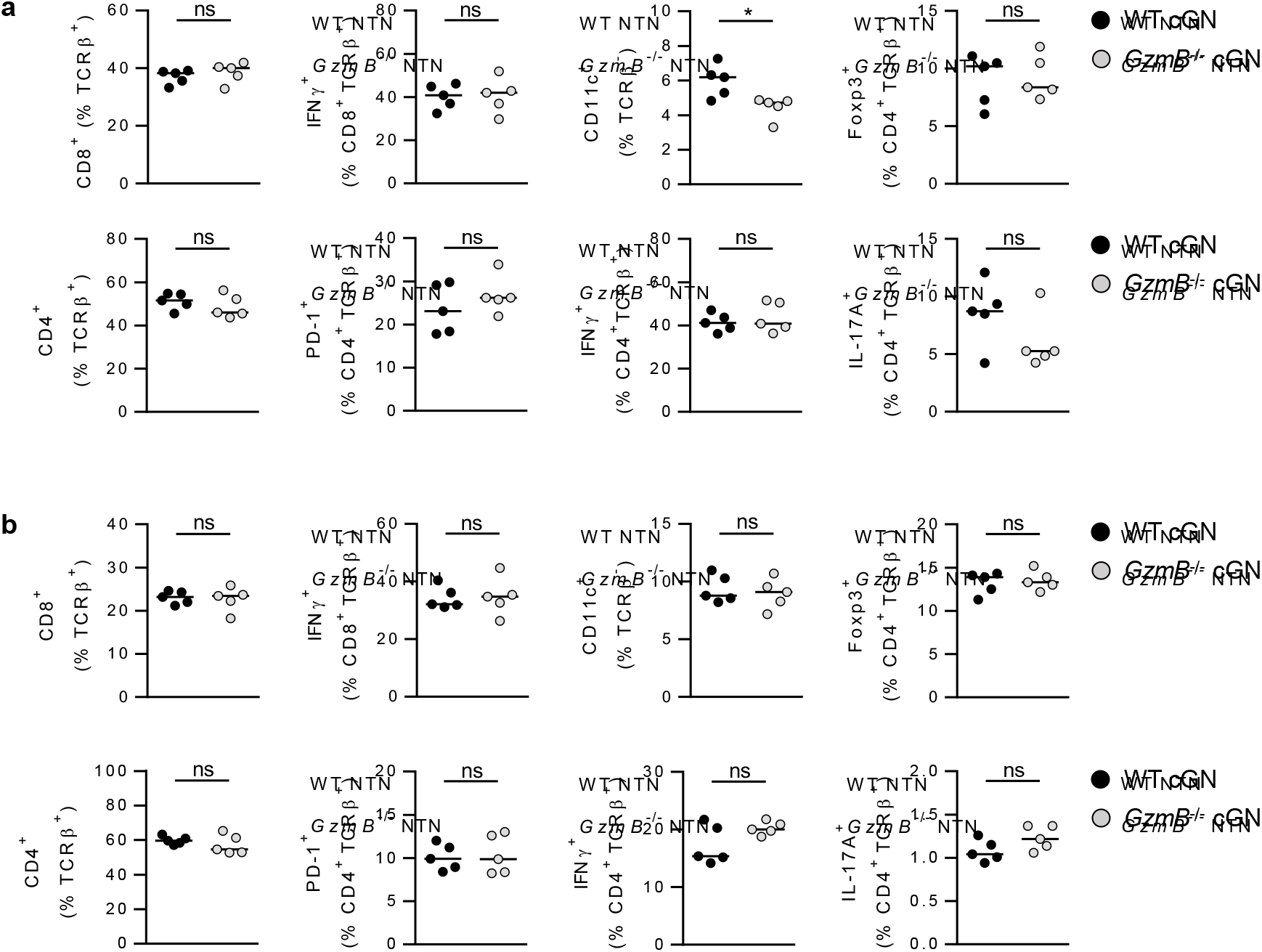
Renal and splenic immune cell populations in nephritic *GzmB*^-/-^ mice. Crescentic GN was induced in *GzmB*^-/-^ and WT mice, which were analyzed 8 days later. Frequencies and phenotypes of (**a**) renal and (**b**) splenic CD8^+^ T cells, CD11c^+^ DCs, Foxp3^+^ Tregs and CD4^+^ T cells were analyzed. Medians of one out of two experiments are shown. ns: not significant.

